# Genotypic and phenotypic diversity of *Staphylococcus aureus* from cystic fibrosis lung infections and their interactions with *Pseudomonas aeruginosa*

**DOI:** 10.1101/814152

**Authors:** Eryn E. Bernardy, Robert A. Petit, Vishnu Raghuram, Ashley M. Alexander, Timothy D. Read, Joanna B. Goldberg

## Abstract

*Pseudomonas aeruginosa* and *Staphylococcus aureus* are the most common bacteria that infect the respiratory tract of individuals with the genetic disease cystic fibrosis (CF); in fact, *S. aureus* has recently overtaken *P. aeruginosa* to become the most common. Substantial research has been performed on the epidemiology of *S. aureus* in CF; however, there appears to be a gap in knowledge in regard to the pathogenesis of *S. aureus* in the context of CF lung infections. Most studies have focused on a few *S. aureus* isolates, often exclusively laboratory adapted strains, and how they are killed by *P. aeruginosa*. Because of this, little is known about the diversity of *S. aureus* CF lung isolates in both virulence and interaction with *P. aeruginosa*. To begin to address this gap in knowledge, we recently sequenced 65 clinical *S. aureus* isolates from the Emory CF Biospecimen Registry and Boston Children’s Hospital, including the reference isolate JE2, a USA300 strain. Here, we analyzed antibiotic resistance genotypes, sequence type, clonal complex, *spa* type, and *agr* type of these isolates. We hypothesized that major virulence phenotypes of *S. aureus* that may be associated with CF lung infections, namely toxin production and mucoid phenotype, would be retained in these isolates. To test our hypothesis, we plated on specific agars and found that most isolates can hemolyze both rabbit and sheep blood (67.7%) and produce polysaccharide (69.2%), consistent with virulence retention in CF lung isolates. We also identified three distinct phenotypic groups of *S. aureus* based on their survival in the presence of nonmucoid *P. aeruginosa* laboratory strain PAO1 and its mucoid derivative. Altogether, our work provides greater insight into the diversity of *S. aureus* CF isolates, specifically the distribution of important virulence factors and their interaction with *P. aeruginosa*, all of which have implications in patient health.

**Author Summary:** *Staphylococcus aureus* is now the most frequently detected pathogen in the lungs of individuals who have cystic fibrosis (CF), followed closely by *Pseudomonas aeruginosa*. When these two pathogens are found to coinfect the CF lung, patients have a significantly worse prognosis. While *P. aeruginosa* has been rigorously studied in the context of bacterial pathogenesis in CF, less is known about *S. aureus*. Here we present an in-depth study of 64 *S. aureus* CF clinical isolates where we investigated genetic diversity utilizing whole genome sequencing, virulence phenotypes, and interactions with *P. aeruginosa*. We have found that *S. aureus* isolated from the CF lung are phylogenetically diverse, most retain known virulence factors, and they vary in interactions with *P. aeruginosa* from highly sensitive to completely tolerant. Deepening our understanding of how *S. aureus* responds to its environment and other microbes in the CF lung will enable future development of effective treatments and preventative measures against these formidable infections.

## Introduction

Cystic fibrosis (CF) is an inherited genetic disease that affects over 100,000 people worldwide [1] and is characterized by a mutation in the **<u>c</u>**ystic **<u>f</u>**ibrosis **<u>t</u>**ransmembrane conductance **<u>r</u>**egulator (CFTR). When CFTR function is compromised, mucus accumulates in the respiratory tract creating a breeding ground for chronic bacterial lung infections. These difficult to treat infections are the predominant cause of morbidity and mortality for people with CF [2].

*Staphylococcus aureus* and *Pseudomonas aeruginosa* are the two most common bacterial species associated with chronic lung infections in cystic fibrosis (CF), with *S. aureus* having recently overtaken *P. aeruginosa* as the most frequently detected bacterial pathogen in sputum samples from all CF patients [1]. In fact, as of 2016, according to the Cystic Fibrosis Foundation’s annual report, 71% of all individuals with CF were infected with *S. aureus*, and 26% were infected with methicillin-resistant *S. aureus* (MRSA) [1]. Historically, *S. aureus* was isolated in younger CF patients and then with age was replaced with *P. aeruginosa* as the dominant species. However, there are a significant number of patients that are coinfected with both *S. aureus* and *P. aeruginosa* [3]. A number of studies [4, 5], including those from our group [6], have shown that coinfection is associated with diminished lung function and more rapid pulmonary decline. The mechanisms responsible for this worsening of disease severity is a topic of intense interest [3, 7, 8], however exactly what promotes this decreased lung function is not known. Research attempting to understand this health decline has mainly focused on the killing interactions between these two microbes, where it has been noted that *P. aeruginosa* readily kills *S. aureus in vitro* [9-12]. Furthermore, we have previously found that the mucoid phenotype of *P. aeruginosa*, which is associated with a chronic infection state, aids in coexistence with *S. aureus* [13], but this was performed on one reference isolate of *S. aureus* from wound infection, JE2.

To date, the importance of *S. aureus* in CF remains controversial [8], as *P. aeruginosa* has historically been recognized as the major pathogen. Perhaps for this reason, the majority of studies on the pathogenesis of *S. aureus* in CF have focused on its interaction with *P. aeruginosa* in the context of coinfection. While this is important, understanding the diversity of *S. aureus* CF isolates themselves as well as their interactions with *P. aeruginosa* remains understudied. For instance, there have been relatively few large-scale comparative whole genomic sequence data analyses using *S. aureus* isolates from patients with CF and none that have observed the corresponding interaction with *P. aeruginosa* [14-20].

*P. aeruginosa* and *S. aureus* are formidable pathogens that are known to alter their virulence phenotypes when shifting from acute to chronic CF lung infection. Substantial research on *P. aeruginosa* has shown significant changes in virulence phenotypes after chronic coinfection in the CF lung, most notably are changes in extracellular products [21]. Less is known about *S. aureus* phenotypic adaptations during chronic infection. Chronic *S. aureus* infection has been characterized by development of small colony variants and a mucoid phenotype, when the bacteria overproduces polysaccharide, both which aid in persistence [22-28]. However, the prevalence of these phenotypic changes in a large number of *S. aureus* CF clinical isolates is not well documented. Moreover, many *S. aureus* virulence factors are toxins [29], and studies investigating toxin production or function in a large number of CF clinical isolates have not been performed.

Our results presented here deepen the current understanding of diversity across *S. aureus* isolates infecting individuals with CF as well as their interactions with *P. aeruginosa*. We obtained 64 clinical isolates of *S. aureus* from individuals with CF from both the Cystic Fibrosis Biospecimen Registry (CFBR; a part of the Children’s Healthcare of Atlanta and Emory University Pediatric CF Discovery Core) and Boston Children’s Hospital. Isolates were chosen to obtain a breadth of patient age, MRSA status, and whether or not these isolates were coinfected with other organisms as detected by the clinical microbiology laboratory. Here, we chose to investigate genotypes and phenotypes believed to be important for *S. aureus* infection in the CF lung, namely Staphylococcal protein A (*spa*) and accessory gene regulator (*agr*) type, antibiotic resistance genes, hemolysis, polysaccharide production, and interaction with *P. aeruginosa*. We hypothesized that *S. aureus* isolates from CF lung infection would be unique from other *S. aureus* isolates based on genome sequence, and that these isolates may lose virulence phenotypes based on what we know of *P. aeruginosa* adaptation in the CF lung. Although our hypotheses were ultimately rejected, we increased our understanding of *S. aureus* in CF. The strains came from genotypes mostly common to US healthcare settings rather than a CF-specific clade, and most *S. aureus* isolates retained virulence-associated genotypes and phenotypes, although a small number seemed incapable of hemolysis or polysaccharide production suggesting possible adaptation to the lung as predicted. Also, not all *S. aureus* isolates behaved the same in regard to interaction with *P. aeruginosa*, signifying the importance to continue studying this interaction. Together, these studies show the large diversity of *S. aureus* isolates infecting individuals with CF, and how this diversity should be further investigated to better inform the treatment of these infections.

## Results

### *Genomic characterization of* S. aureus *CF clinical isolates including antibiotic resistance and virulence genes*

To begin to determine the diversity of *S. aureus* isolates from individuals with CF, we previously reported the genome sequences of 64 *S. aureus* isolates collected from 50 individuals with CF and the reference isolate JE2 (total of 65 isolates) as described in Bernardy, et al 2019 [30]. *S. aureus* JE2, a derivative of USA300, was used throughout our study as a control non-CF associated strain because its sequence and phenotypes were known [31-33]. Our CF clinical isolates were obtained from patients with a wide range of ages and were from two different sites (CFBR and Boston Children’s Hospital, Table 1). We first created a phylogeny to indicate how similar our isolates were to one another. Fig 1 shows the isolates analyzed here represent 8 phylogenetically diverse clonal complexes (CC) of the 66 we defined previously within the > 40,000 *S. aureus* genomes compiled to date in the publicly available Staphopia database [34]. The most common clonal complexes (CC) represented in these isolates were CC5 and CC8, which are also the most prevalent hospital-acquired **m**ethicillin-**r**esistant ***S. a****ureus* (MRSA) in the USA [35]. In fact, forty of the 64 clinical isolates (all CC5 or CC8) were MRSA. The **m**ethicillin-**s**ensitive ***S. a****ureus* (MSSA) isolates were more genetically diverse and included some CC5 and CC8 and one CC398 (livestock strain).

**Table 1.**
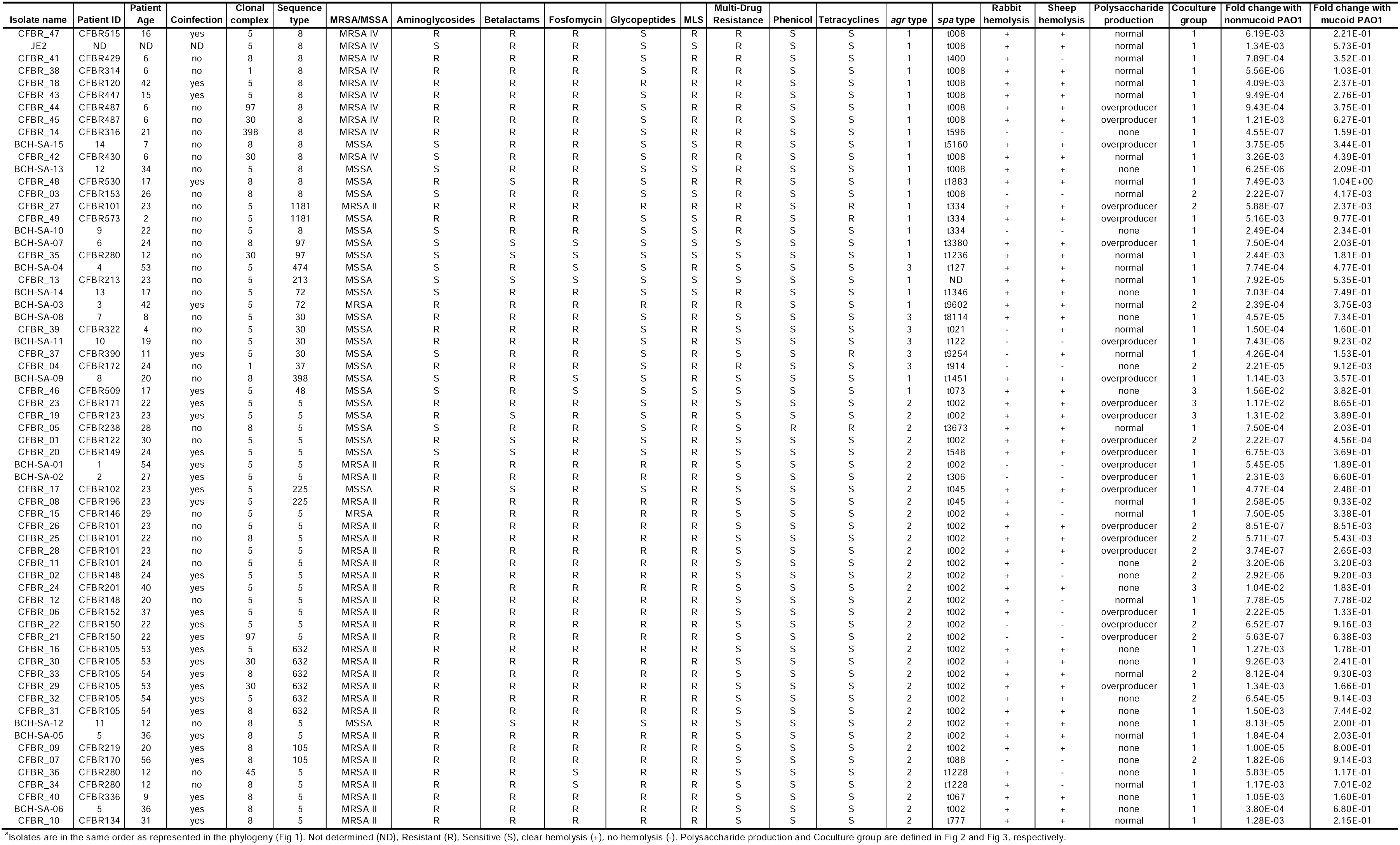
Compilation of metadata, genotypes and phenotypes from CF clinical isolates of *S. aureus* and laboratory strain JE2.^**a**^.

**Fig 1.**
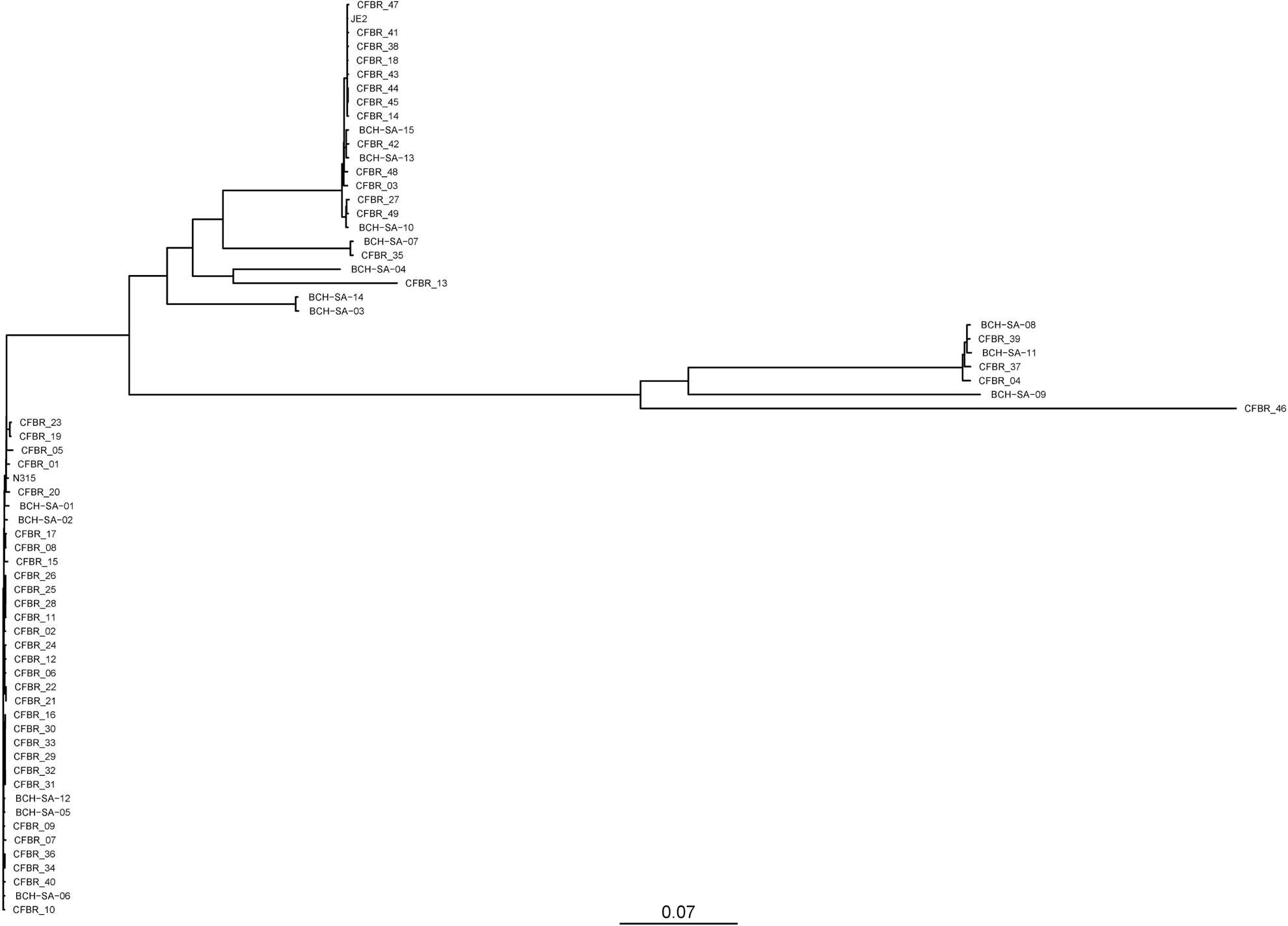
Whole genome phylogeny. Maximum-likelihood phylogenetic tree showing evolutionary relationships between the core genomes of *S. aureus* reference genome N315 and 65 isolates from this study.

We next analyzed the sequences of all 65 isolates (64 clinical isolates and JE2) for 15 antibiotic resistance genes, *agr* type, *spa* type, and alpha and beta toxin genes. The most prevalent antibiotic resistance genes in our set of isolates include fosfomycin, ß-lactams, MLS (macrolides, lincosamides, streptogramines), and aminoglycosides present in 57, 56, 54, and 49 isolates, respectively (Table 1). No isolates had resistance genes to fusidic acid, rifampin, sulfonamides, thiostrepton, trimethoprim, and tunicamycin. Most isolates (33 out of 65) had 6 resistance genes, while JE2 had 4. The most resistance genes present in one isolate was 8, and only three isolates had no resistance genes.

The accessory gene regulatory (*agr*) quorum sensing system controls multiple virulence factors in *S. aureus* and is thought to be vital in infection [36-38]. There are four known types of this system based on mutations and polymorphisms in the histidine kinase and autoinducer peptides [39]. Among our isolates, there were 24 *agr* type I, 35 type II, and 6 type III (Table 1). JE2 was known as *agr* type I, which was confirmed by our sequence analysis. None of our isolates were *agr* type IV; this reflects the phylogeny of the strains and what is known about *agr* type IV due to its main association with isolates from skin infections [40].

Staphylococcus protein A (*spa*) is implicated in virulence and typing of this gene by identifying the specific repeats in its variable repeat region has historically been used to distinguish between circulating variants of *S. aureus* during an outbreak. Among our isolates, the two most common *spa* types are t002 (25 isolates) and t008 (10 isolates) (Table 1), both of which are common types in the United States [41]. JE2 was confirmed as *spa* type t008 in our analysis. None of our isolates appear to be a part of an outbreak based on date of collection, *spa* type, clonal complex (CC), and sequence type (ST), meaning no isolates collected around the same time shared genotypes typically used for classification.

Finally, we wanted to determine if each isolate had alpha and beta toxin genes (*hla* and *hlb*, respectively), two important toxins involved in infection [29, 42, 43]. Sequences from hemolysis positive JE2 were used for comparison. All 64 clinical isolates had both genes in their genomes. Therefore, we checked the amino acid sequence identity with a known toxin producer (JE2) in order to infer function. Only 14 isolates had 100% identity with alpha toxin, but another 43 had at least 99% identity. For beta toxin, 21 out of 65 isolates had 100% identity with a functioning beta toxin, and 44 had at least 99% identity. We are currently investigating these sequences for causative mutations for those isolates lacking hemolysis.

### S. aureus *CF isolates hemolyze blood and produce polysaccharide*

*S. aureus* utilizes an arsenal of toxins as virulence factors during infection [29]. While detecting toxin genes in *S. aureus* CF clinical isolates is commonly performed in epidemiology studies [44], the prevalence of toxin production in a large set of *S. aureus* CF clinical isolates has not been performed. Therefore, in order to assess the virulence capabilities of our *S. aureus* clinical isolates, we measured the production of hemolysins on blood agar plates. Two of the most prominent hemolytic toxins are alpha and beta toxin, whose production can be tested by observing clear hemolysis on rabbit and sheep blood agar plates, respectively [45, 46]. Of the 65 isolates tested, 44 hemolyze both rabbit and sheep blood (Rabbit + / Sheep +, Table 2), confirming alpha and beta toxin production, 10 could not hemolyze either blood agar (Rabbit -/ Sheep -, Table 2). The remaining 11 isolates were positive for only one type of blood hemolysis (Rabbit + / Sheep - or Rabbit -/ Sheep +, Table 2). Even though all isolates had both alpha and beta toxin genes present, the activity is not apparent in some of these isolates likely due to mutations elsewhere in the genome.

**Table 2.**
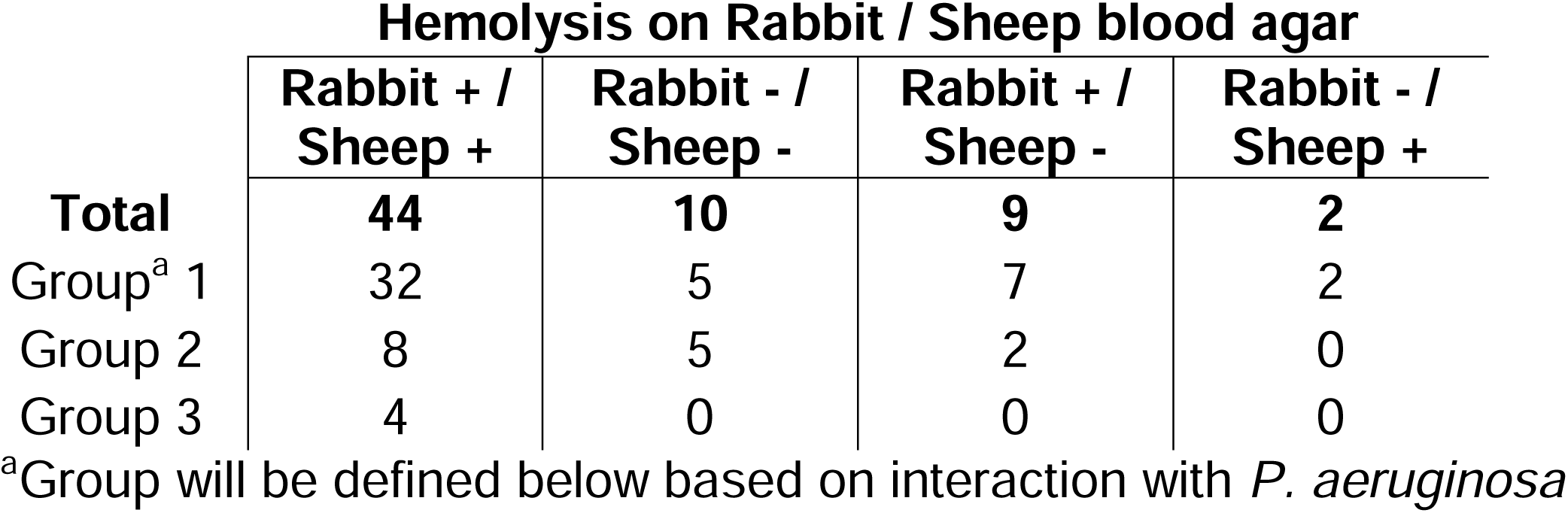
*S. aureus* CF isolates positive (+) or negative (-) for hemolysis on blood agar plates.

Another important virulence factor for *S. aureus* is the mucoid phenotype characterized by an overproduction of the polysaccharide poly-*N*-acetyl-β-(1,6)-glucosamine (PNAG) [47-49]. Therefore, the polysaccharide production of each strain was assessed by streaking onto Congo Red Agar (CRA) plates as described previously [47, 50]. Results were interpreted by observing both color and appearance (smooth vs. rough) of colonies on plates. Genetically defined isogenic mutants of MN8 served as controls [49]. We observed three different phenotypes and an example of each of these is plated on CRA in Fig 2. Among the 65 isolates tested, there was an even split among non-producers (20 out of 65), normal polysaccharide producers (23 out of 65), and overproducers (22 out of 65) (Table 1). Overall, 45 isolates (69.2%) were capable of producing polysaccharides to some degree, suggesting that this phenotype is conserved in *S. aureus* CF clinical isolates.

**Fig 2.**
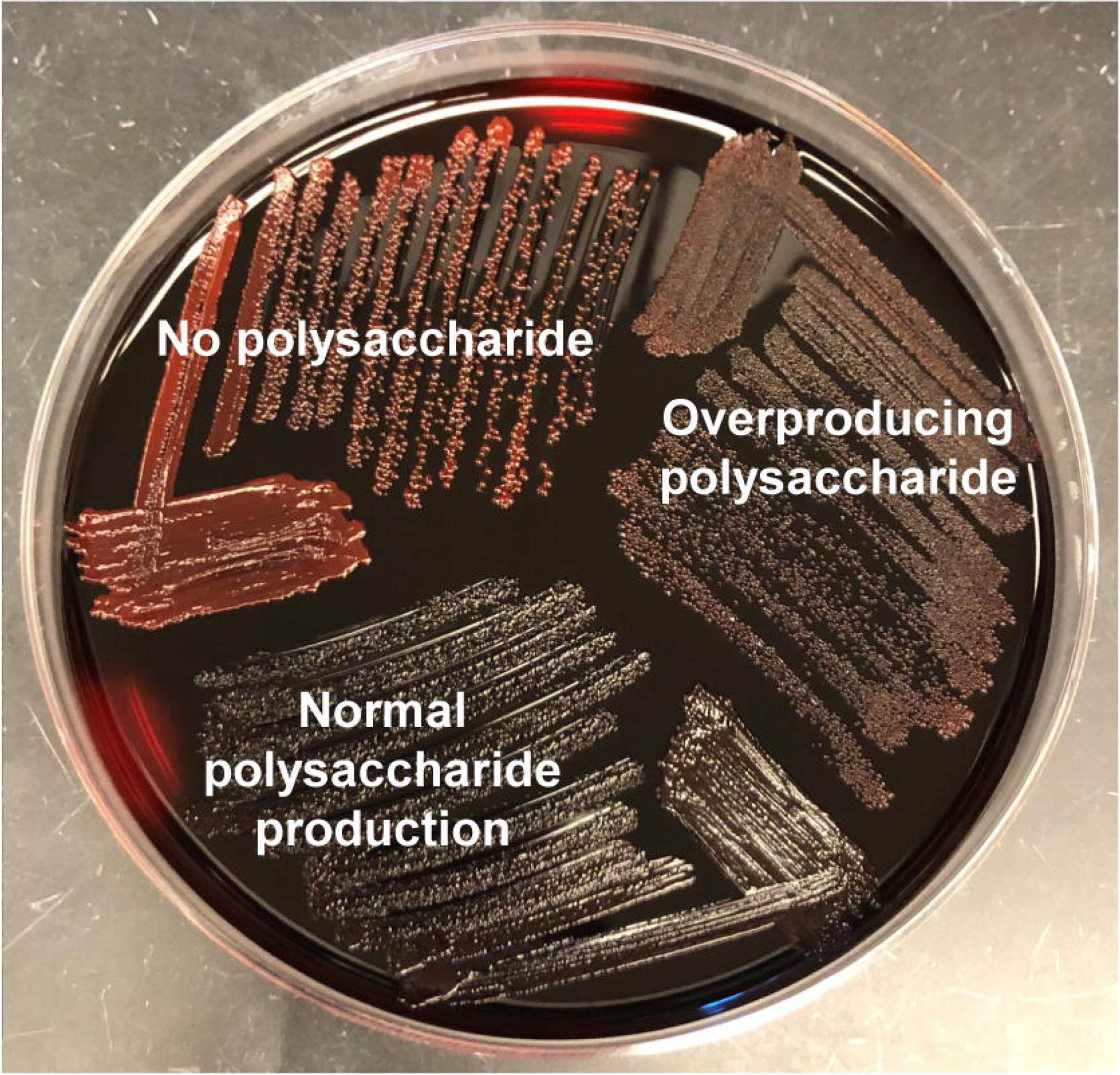
Plating on Congo Red Agar shows 3 phenotypes. One representative isolate of each phenotype is shown. Isolates classified as having “no polysaccharide” are bright red in color and smooth or shiny in appearance. Isolates classified as having “normal polysaccharide production” are much darker in color than those with “no polysaccharide”, many were black in color as shown in this figure. These isolates are also smooth or shiny in appearance. Finally, isolates classified as having “overproducing polysaccharide” are also darker than “no polysaccharide” but have a rough or matte appearance.

The *ica* operon is known to be responsible for this phenotype, specifically a 5bp deletion upstream of the *icaA* gene is known to confer a mucoid or overproducer phenotype [47, 49]. Interestingly, none of these isolates had this deletion, indicating the presence of other mutations that cause this phenotype in these isolates.

### S. aureus *CF isolates fall into three distinct groups based on interactions with P. aeruginosa*

Previously, it has been shown that nonmucoid PAO1 kills *S. aureus* lab isolate JE2, while mucoid PAO1 does not [13]. Therefore, we were curious whether this trend would be maintained with CF clinical isolates of *S. aureus* and consequently all of our 65 *S. aureus* isolates were assessed in a coculture assay with both nonmucoid and mucoid PAO1. While a number of different techniques have been used to look at how *P. aeruginosa* and *S. aureus* survive in coculture [13, 51-55], we have developed a simple, repeatable, well-controlled, quantitative assay for monitoring these interactions *in vitro*. This coculture assay allows for these bacteria to come in contact with one another on a solid surface as they might in a biofilm in the lung. We are also able to grow each species by itself under the same conditions in order to better understand how the bacterial survival changes when grown in coculture versus alone.

To determine whether or not each *S. aureus* isolate is killed in the presence of *P. aeruginosa*, fold change of *S. aureus* grown in coculture with *P. aeruginosa* compared to when it grew alone in the same conditions was calculated (Fig 3). A fold change of less than 10^−2^ (or >100 fold decrease in cfu/ml when grown with *P. aeruginosa*) was considered significantly killed (Fig 3, horizontal black lines), based on other coculture assays that measure killing [56]. This definition allowed us to assign each isolate into an interaction group based on how it interacted with nonmucoid PAO1 and mucoid PAO1. Group 1 isolates are those that fit with the previous trend observed and are only killed by nonmucoid PAO1 (Fig 3A). We observed a range in fold changes, but each fit this same trend; the black bars designating “fold change when grown with nonmucoid PAO1” are all below the killing line, while the grey bars designating “fold change when grown with mucoid PAO1” are all above the killing line. Most isolates fit in this group (46 out of 65, Fig 3A), including previously tested isolate JE2. Group 2 isolates were those killed by both nonmucoid and mucoid PAO1 (15 out of 65, Fig 3B) where both black and grey bars are below the killing line. Finally, group 3 isolates were those killed by neither PAO1 strain (4 out of 65, Fig 3C) where both bars are above the killing line.

**Fig 3.**
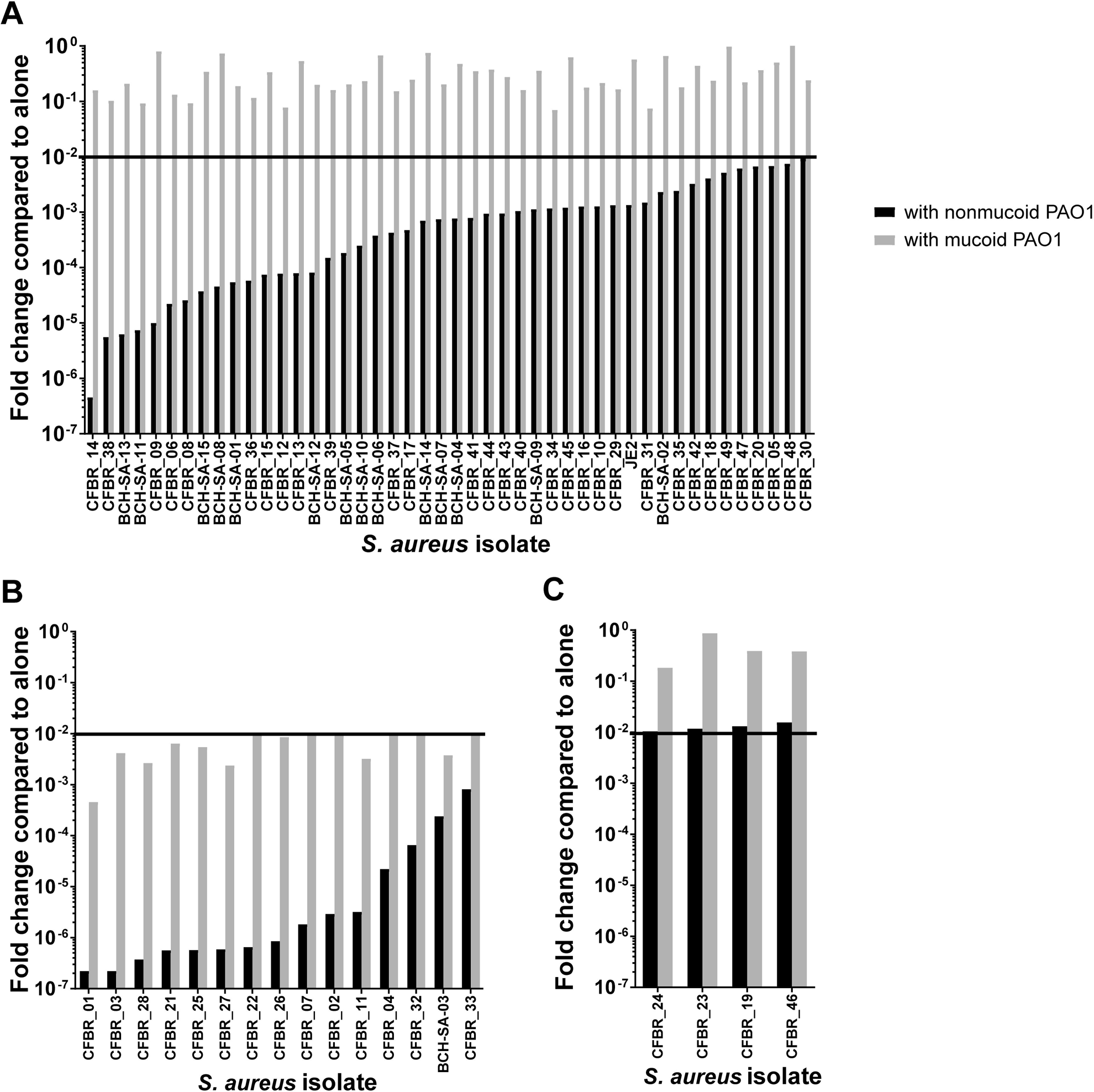
Coculturing *S. aureus* CF isolates with nonmucoid and mucoid PAO1 revealed 3 interaction groups. Fold change of each *S. aureus* isolate after coculture with both nonmucoid and mucoid PAO1. Fold change was calculated by dividing the cfu/ml of *S. aureus* grown with nonmucoid PAO1 (black bars) or with mucoid PAO1 (grey bars) over the cfu/ml of each *S. aureus* isolate grown alone. Fold change of <10^−2^ was considered to be significantly killed as represented by the horizontal black line, denoted as “killing line” in text. (A) Group 1 isolates, those killed by nonmucoid PAO1 only, have black bars below 10^−2^ while all grey bars are above this threshold. There were 46 isolates that fit into this interaction group, including the previously tested lab isolate JE2. (B) Group 2 isolates, those killed by both nonmucoid and mucoid PAO1, have both black and grey bars below the 10^−2^ threshold. 15 isolates are in this interaction group. (C) Group 3 isolates, those killed by neither PAO1 strain, have both black and grey bars above the 10^−2^ threshold. 4 isolates are in this group. The average of technical triplicate of one experiment representative of the three biological replicates performed are shown.

None of the *S. aureus* isolates that we tested affected growth of either the nonmucoid or the mucoid PAO1 strain in this assay (S1 Fig). We also determined that the decrease in survival when *S. aureus* is grown with *P. aeruginosa* is due to killing not growth inhibition. This was shown by performing the same coculture assay, but instead of taking one time point at 24 hours, we measured cfu/ml of *S. aureus* alone and with *P. aeruginosa* at multiple time points. We observed that *S. aureus* grows well with *P. aeruginosa* until approximately 12 hours when its cell numbers drop dramatically (S2 Fig), suggesting that the dramatic fold changes we see are, in fact, killing and not due to growth inhibition. Finally, these *S. aureus* isolates did not inherently have different growth patterns. As shown in S3 Fig, all *S. aureus* isolates when grown alone grow to approximately the same cfu/ml (∼10^9^) over 24 hours in the coculture assay conditions. Therefore, we conclude that the differences in survival of the various *S. aureus* isolates when grown with *P. aeruginosa* are not due to inherent differences in growth.

## Discussion

### Phylogeny relatedness to genotypes and phenotypes

To understand the diversity present in this set of 64 *S. aureus* CF clinical isolates and the reference isolate JE2 (65 total), we sought to combine our genotypic and phenotypic data as well as metadata obtained from the clinical lab. Most isolates belonged to CC5 and CC8 which is consistent with the most commonly acquired hospital MRSA isolates, possibly suggesting nosocomial acquisition. In Table 1, the isolates are in the order placed in the phylogeny in Fig 1. Therefore, we can observe genotypes and phenotypes that appear linked based on full genome sequence relatedness. As expected, clonal complexes and sequence types cluster together, as well as the MRSA/MSSA phenotype, *agr* type, and *spa* type (Table 1).

When comparing phylogenetic relatedness and observed phenotypes, we revealed some interesting connections. While rabbit and sheep hemolysis do not seem to obey any order based on relatedness, comparing the polysaccharide production phenotypes in Table 1, there are more normal polysaccharide-producing isolates grouped together, followed by mostly overproducers, then mostly nonproducers. This clustering of phenotypes suggests a genetic association; however, we know these differences are not due to the 5bp deletion in the *ica* operon, as discussed previously. Coinfection has a similar pattern with isolates not from coinfection clustering together (Table 1, near the top) and those from coinfection clustering together (Table 1, near the bottom). We plan to continue to use the sequencing data we obtained to determine a genetic basis for these phenotypes.

There was one connection between phylogeny and phenotypes that was notably absent; there was no observed relationship between coculture interaction group and phylogeny. As seen in Table 1, there is an instance where groups cluster together, specifically, 3 out of 4 Group 3 isolates cluster together. But Group 1 and 2 isolates appear well distributed throughout the phylogeny. This suggests that what makes these isolates a specific coculture group is more complex than originally anticipated. We are currently further investigating these sequences to determine genetic factors involved in the interaction between *P. aeruginosa* and *S. aureus*.

### Longitudinal isolates

Some of our isolates were longitudinal; they came from the same individual with CF (Patient ID column, Table 1) and were collected over a period of time. In both Table 1 and Fig 1, the longitudinal isolates cluster together on the phylogeny, suggesting that these isolates all originated from one initial infecting isolate, or at least isolates that are very closely related. It is also interesting, although not unexpected, to see that many of their phenotypes were similar. For example, most isolates collected from the same individual had the same hemolysis abilities (Table 1, Rabbit hemolysis and Sheep hemolysis columns). However, one patient, CFBR105, provided four isolates in Group 1, and two isolates in Group 2. When arranged by date of collection, the Group 2 isolates were both collected later, while all having the same ST, suggesting that these isolates may have evolved inside the lung to no longer defend against mucoid *P. aeruginosa* attack. These isolates also have a range of polysaccharide production, suggesting more adaptation over time. However, an alternate hypothesis is that these isolates were always present in the lung growing in a population, but during collection only a single colony was chosen. We are currently further investigating additional isolates collected from the same individuals to help us determine the genetic factors involved in adaptation to the CF lung, as well as diversity of the infecting *S. aureus* populations.

### Hemolysis and polysaccharide production were common among our isolates

Toxins are an important part of *S. aureus* virulence, and alpha toxin especially has been shown to be required for pulmonary infection in CF mice [57]. Consistent with this observation, most of the *S. aureus* clinical isolates tested produced hemolytic toxins, determined by their ability to completely hemolyze both rabbit and sheep blood. Interestingly, only 10 isolates (15.4%) were unable to hemolyze either blood agar. The ubiquity of hemolytic activity in these isolates supports the knowledge that *S. aureus* toxins are important when infecting the human host [29].

*S. aureus* polysaccharide production has been implicated to be important for chronic colonization in the CF lung [47]. Qualitative phenotypic characterization of polysaccharide production following plating on CRA in this study shows that consistent with this hypothesis, most isolates were capable of producing polysaccharide (both normal producers and overproducers). Furthermore, the fact that none of the CF isolates had the conserved 5bp deletion upstream of the *icaA* gene known to confer a mucoid or overproducer phenotype [47, 49], suggests a different mechanism of polysaccharide production, which we are currently investigating. Twenty isolates (30.8%) were characterized as non-producers due to their red color on CRA plates. These isolates may have other mechanisms for attachment and biofilm production outside of this specific polysaccharide or may benefit in some other way by not adhering to surfaces. We are currently investigating the genomes of these isolates for the cause of this lack of polysaccharide production. Overall, polysaccharide production was common among these clinical isolates, suggesting that this behavior is important in CF lung infection.

### Group 1 isolates may come from initial infection

While it is generally understood that *P. aeruginosa* kills *S. aureus in vitro* [9-13, 51, 58] and is hypothesized to also do so *in vivo*, these studies were performed on a small number of strains and focused on how *P. aeruginosa* is killing *S. aureus*. Our studies here allow us to determine if *P. aeruginosa* killing *S. aureus* is a phenotype typical of CF isolates. Our group previously showed that JE2, an isolate from wound infection, was killed by nonmucoid PAO1 but survived when cocultured with mucoid PAO1, and subsequently discussed the mechanism behind this conclusion [13]; therefore, we examined if CF isolates behaved similarly. The majority of our isolates (46 of 65), including JE2, were killed by nonmucoid PAO1 only and we subsequently called these our Group 1 isolates. While it is not surprising that these isolates behaved this way due to previously defined mechanisms in Limoli, et al. [13], it is interesting that isolates from lung infection behave the same as an isolate from a wound infection in regards to interaction with *P. aeruginosa*. Remarkably, when looking at Group 1 isolates in Table 1, many were from younger patients and were sensitive to aminoglycosides and glycopeptides. This observation suggests that these isolates may be from the initial stages of infection. There is a known switch in predominance of infection from *S. aureus* in childhood to *P. aeruginosa* in adults with CF which aligns with our observation, because *S. aureus* isolates from initial infection could be sensitive to nonmucoid *P. aeruginosa* which would allow for this switch to occur.

### Group 2 isolates were incapable of hemolysis, but more resistant to antibiotics

While most isolates were in Group 1, we recognized a set of isolates that were killed by both nonmucoid and mucoid PAO1, denoted Group 2 (15 out of 65 isolates). This observed phenotype was surprising to us based on how often these two bacteria are thought to coinfect the CF lung. It is likely that these isolates have not come in direct contact with *P. aeruginosa* and therefore have not had a need to develop defensive strategies. These isolates may also have other mutations that increased their fitness in the CF lung that coincidentally led to them being less competitive with *P. aeruginosa*. In line with this hypothesis, half of the isolates incapable of hemolyzing either type of blood agar (Rabbit -/ Sheep -, Table 2) were in coculture Group 2. Therefore, it is tempting to speculate that these isolates may have lost some virulence phenotypes resulting in lack of hemolysis and increased susceptibility to killing by *P. aeruginosa*. While looking at Group 2 isolates in Table 1, we noticed that many were from older patients, and were resistant to methicillin (MRSA), aminoglycosides, and glycopeptides. These observations may suggest that these isolates come from chronic infection due to the increase in antibiotic resistances and age of the patient from which they were collected. There is also research suggesting an inverse relationship between toxin production and ability to cause infections, with low-cytotoxic isolates causing more pervasive infections [59]. This data supports our hypothesis that these isolates might cause chronic infection because many Group 2 isolates were negative for hemolysis. As previously mentioned, some of our longitudinal isolates switched to a Group 2 over time in the same patient, consistent with this data. Therefore, we might be observing a *S. aureus* adaptation over time in the CF lung where they lose expression of virulence factors, similarly to *P. aeruginosa*. It is interesting to consider that during CF lung infection, it might be more advantageous for *S. aureus* to retain antibiotic resistance phenotypes rather than relinquish them in favor of coexistence with *P. aeruginosa*, leading to a Group 2 interaction phenotype.

### Group 3 isolates were coinfected with P. aeruginosa *at time of collection*

The most surprising group of isolates are those that were resistant to killing by both nonmucoid and mucoid PAO1, denoted Group 3 (4 of 65 isolates). *P. aeruginosa* is a potent competitor and utilizes an arsenal of extracellular products and other mechanisms to kill neighboring bacteria; therefore, it is interesting that these *S. aureus* isolates have developed defensive strategies to survive coculture *in vitro*. Unsurprisingly, when investigating the metadata associated with these isolates, we discovered that all four of these isolates, which each came from a different individual, were coinfected with *P. aeruginosa* at the time of collection. Of the 46 Group 1 isolates, 20 (43.4%) were coinfected with *P. aeruginosa* at the time of collection, while 7 of the 15 (46.7%) Group 2 isolates were from coinfected individuals, signifying that coinfection was most important for Group 3 isolates. While much work has been done to show that *P. aeruginosa* and *S. aureus* do not appear to come in direct contact during an established infection in a chronic wound model [60, 61], whether this holds true in the context of CF has not been clearly shown. In CF, it is possible that during initial infection there is interaction between these two microbes, but that they eventually separate and create spatial structure due to their antagonistic interaction, following a well-studied ecological theory [62]. It is also possible that these *S. aureus* isolates obtained other fitness benefits from genetic changes that allow coexistence with *P. aeruginosa*.

### Diversity of S. aureus *CF isolates*

Based on the analysis performed here, we conclude that *S. aureus* CF clinical isolates are more diverse than we hypothesized. Not only do they vary in virulence phenotypes but also in their interactions with *P. aeruginosa*. These variations may be due to inherent differences during initial infection, or evolutionary changes in response to their environment, both that of the CF lung but also of the presence of other pathogens like *P. aeruginosa* or other members of the lung microbiome [63]. Most isolates retained virulence-associated phenotypes, namely hemolytic activity and mucoid phenotype, after infecting the CF lung. The mucoid phenotype may aid in adhesion and can protect *S. aureus* from immune cell attack, so it was not surprising to find that many isolates were mucoid. Many studies had previously shown that *P. aeruginosa* kills *S. aureus* in a variety of *in vitro* experiments. Some have also shown *P. aeruginosa* isolates that cannot kill *S. aureus*, however a widespread examination of *S. aureus* isolates and their ability to withstand *P. aeruginosa* attack had not been performed before this study. We are excited to report the amount of diversity in this interaction and plan on further investigating the genetic factors involved by utilizing various molecular techniques and the sequencing data we have obtained.

While we have not yet identified the *S. aureus* mechanisms involved in the diversity of interaction with *P. aeruginosa*, we have ruled out some specific genotypes and phenotypes. We conclude that this interaction is complex and multifactorial. There were no striking phenotypes or genotypes that were specific or unique for each coculture interaction group. For example, *S. aureus* polysaccharide PNAG has been implicated in persistence [47] and we know that polysaccharide produced by *P. aeruginosa* can alter interaction with *S. aureus*. Therefore, we hypothesized that overproduction of *S. aureus* polysaccharide could possibly provide protection from *P. aeruginosa* attack, making these isolates Group 3. However, Group 3 isolates had varying polysaccharide production as determined in this study (Table 1). We also tested the positive and negative polysaccharide producing controls (MN8 WT, mucoid, and Δ*ica*) in our coculture assay and all were killed by nonmucoid PAO1 but not mucoid PAO1, and therefore were Group 1 (data not shown). Consequently, based on our data, we do not believe polysaccharide production by *S. aureus* to be important in this interaction. We are currently further investigating these *S. aureus* isolates for genetic factors or phenotypes responsible for the varying interaction with *P. aeruginosa*.

Interestingly, none of the *S. aureus* isolates discussed in this study were small colony variants, which has been shown to be a known adaptation to the CF lung environment [22-25]. Some became small colony variants after being challenged with *P. aeruginosa*, but it was not consistent, and they quickly reverted when re-streaked alone on SIA (data not shown). This leads us to a limitation of our study; we only collected single isolates from a patient, which may have led us to lose small colony variants which are thought to be the dominant form of *S. aureus* in the CF lung [64]. In future studies, we will obtain multiple colonies from the same patient to better understand the diversity of *S. aureus* inside a single individual. We also only performed coculture tests using nonmucoid and mucoid PAO1 which may not fully represent these isolates true abilities to coexist or antagonize *P. aeruginosa* inside the CF lung. In the future, we hope to expand on this by testing clinical isolates of both *S. aureus* and *P. aeruginosa*.

*S. aureus* is the most prevalent cause of lung infection in individuals with CF, yet a small number of large-scale studies had been performed combining genomic and phenotypic data before this study. Understanding the diversity of these isolates and how specific phenotypes and genes connect to patient health is paramount to better treating these patients. If we can provide clinical microbiology labs with a list of specific *S. aureus* traits to monitor in order to prevent coinfection between *P. aeruginosa* and *S. aureus* and the associated health decline, we could make a huge impact on the health of individuals with CF. Outside of lung infections, MRSA causes a substantial number of infections at all body sites and is recognized as a significant threat by the CDC. We have identified a subset of isolates that are sensitive to attack by other bacteria. If we can identify what kills these bacteria, or what genes make them sensitive, it could provide new treatment options for these notoriously hard to treat infections. Overall, our work contributes to a better understanding of the diversity of *S. aureus* in CF lung infections.

## Materials and Methods

### Bacterial strains and growth conditions

The *S. aureus* isolates used in the study are listed in Table 1. *P. aeruginosa* isolates used were laboratory strain nonmucoid PAO1 [65] and mucoid PAO1 containing *mucA22* allele, also known as PDO300 [66]. *P. aeruginosa* and *S. aureus* were grown in lysogeny broth (LB) and trypticase soy broth (TSB), with 1.5% agar for solid medium. Selective media for *P. aeruginosa* was *Pseudomonas* Isolation Agar (PIA, BD Difco), while selective media for *S. aureus* was trypticase soy agar (TSA, BD BBL) with 7.5% NaCl, called *Staphylococcus* Isolation Agar (SIA).

### Whole genome phylogeny

The genomes were processed using the Staphopia Analysis Pipeline [34]. Parsnp [67] and FastTree2 [68] were used for alignment and phylogenetic tree construction.

### Genotypic characterization of virulence phenotypes

Alpha and Beta toxin sequences (*hla* and *hlb* respectively) were extracted from hemolysis positive *S. aureus* JE2 reference genome (Accession: GCA_002085525.1) and were queried against the *S. aureus* CF clinical isolate genomes using BLAST 2.9.0 [69]. A similar strategy was adopted for *agr* typing where AgrD sequences for the 4 *agr* groups were used as references (Accessions: AAB80783.1, AAB63265.1, AAB63268.1, AAG03056.1). Hits with > 95% amino acid sequence identity and >90% query coverage were considered to be positive for presence of alpha/beta toxin or the respective *agr* type.

Staphylococcal protein A (*spa*) type repeat-successions and sequences corresponding to individual repeats were downloaded from the Ridom Spa Server [70]. These files were then combined to create a fasta file having the complete sequence for 18915 *spa* types. These sequences were then converted to a BLAST database and used to query our *S. aureus* genomes. Because *spa* types are assigned based on repeat sequence identity and the number of repeats, we assigned the longest spa sequence having 100% sequence match and 100% query coverage as the *spa* type for a given sample.

### Hemolysis assays

*S. aureus* CF clinical isolates and the lab isolate JE2 were plated on rabbit blood and sheep blood agar plates. Briefly, wooden sticks were placed in cryovial with frozen stock of the chosen strain, then gently touched to the surface of the chosen blood agar plate. Plates were incubated at 37°C for 24 hours. The diameter of each colony size and size of clear hemolysis, if any, was recorded. After which, plates were incubated again at 4°C for 24 hours and colony size and clear hemolysis size were recorded again. For ease, each isolate was scored as “+” if clear hemolysis was detected or “-” for no clearing (not hemolyzing blood). All isolates tested grew on both types of plates.

### Phenotypic characterization of polysaccharide production

Each *S. aureus* CF clinical isolate, along with positive (MN8 wild-type, MN8 mucoid which had a 5bp deletion in *ica* operon) and negative (MN8 Δ*ica* which had the entire *ica* operon deleted) controls for polysaccharide production (provided by Dr. Gerald B. Pier, Brigham & Women’s Hospital, Harvard Medical School) [49] were streaked on Congo red agar plates (CRA). CRA was made as previously described [50]. We combined 18.5g Oxoid brain heart infusion broth, 25g sucrose, and 5g agar in 500ml of distilled water and autoclaved it. After cooling to ∼55°C, we then added 8mLs of Congo Red dye stock solution (5g in 100mL and autoclaved). To streak onto CRA, wooden sticks were placed in cryovials with frozen stock of each strain then streaked across CRA. Plates were incubated at 37°C for 24 hours. Results were interpreted as previously described [47, 71-73]: Black and deep red smooth colonies were considered to be normal polysaccharide producing strains (like MN8 wild-type), red smooth colonies were considered to be non-producers (like MN8 Δ*ica*), and rough colonies of any color (like MN8 mucoid) were considered to be overproducers.

### Coculture assay

To monitor the interaction between *P. aeruginosa* and the *S. aureus* isolates in this study, we performed a quantitative coculture assay using either nonmucoid or mucoid *P. aeruginosa* strain PAO1. Briefly, wooden sticks were placed in cryovials with frozen stocks of nonmucoid and mucoid PAO1 and then streaked for singles onto *Pseudomonas* Isolation Agar (PIA) while *S. aureus* isolates were streaked onto *Staphylococcus* Isolation Agar (SIA). Both were incubated at 37°C overnight. Single colonies were selected and then grown in liquid lysogeny broth (LB) at 37°C overnight. These cultures of *P. aeruginosa* and *S. aureus* were then back-diluted to an optical density (OD) of 0.05 and mixed in a 1:1 ratio with each other, or with sterile broth as “alone” controls. 10 µL of each mixture were placed onto a 0.45 µm filter on a TSA plate and incubated at 37°C for 24 hours. After incubation, filters were removed using sterile forceps and bacteria resuspended in 1.5 mL of sterile LB before serial dilution and plating onto PIA and SIA. After incubation at 37°C overnight, colonies were counted and colony forming units (cfus) calculated. Fold change of *S. aureus* cfu/ml was calculated by dividing the cfu/ml of *S. aureus* grown with nonmucoid PAO1 or with mucoid PAO1 over the cfu/ml of each *S. aureus* isolate grown alone. A fold change of <10^−2^ was considered to be significantly killed.

### Availability of data

Raw Illumina reads available under BioProject accession number PRJNA480016 were used in this study.

## Acknowledgments

This work was supported, in part, by a Pediatric Research Alliance Pilot Project (00068914 to JBG and TR) from Cystic Fibrosis and Airways Disease (CF-AIR) and Children’s Healthcare of Atlanta. Bacterial isolates were obtained from the Cystic Fibrosis Biospecimen Registry, which is supported in part by the CF Discovery Core of the CF@LANTA RDP Center, and by the Center for CF and Airways Disease Research, components of the Emory-Children’s CF Center of Excellence at Emory University and Children’s Healthcare of Atlanta. We would like to thank Dr. Gregory P. Priebe for isolates obtained from the Boston Children’s Hospital, Dr. Gerald B. Pier for *S. aureus* strains for investigating polysaccharide production, and Dr. Cassandra Quave with assistance with the hemolysis assay. We would also like to thank the NIH IRACDA Fellowships in Research and Science Teaching program at Emory for financial support to EEB. (Project Number 5K12GM000680-19). This publication made use of the *spa* typing website (http://www.spaserver.ridom.de/) that is developed by Ridom GmbH and curated by SeqNet.org (http://www.SeqNet.org/).

The content of the manuscript is solely the responsibility of the authors and does not necessarily represent the official views of the funding agencies. The funders had no role in study design, data collection and interpretation, or the decision to submit the work for publication.

## Figure legends

**S1 Fig. No effect on *P. aeruginosa* when cocultured with *S. aureus* for 24 hours.** Average cfu/ml of nonmucoid (top graph, black bars) and mucoid (bottom graph, grey bars) PAO1 after 24 hour coculture with each *S. aureus* isolate. No *S. aureus* isolate tested effected *P. aeruginosa* growth.

**S2 Fig. *S. aureus* Group 1 isolate grown over time with and without PAO1.** Average cfu/ml of a Group 1 *S. aureus* isolate JE2 grown alone (orange), with non-mucoid PAO1 (black), and with mucoid PAO1 (grey) over 24 hours using the same coculture assay procedure as seen in Fig 2. All three experimental groups grew well initially, then the black line (*S. aureus* grown with non-mucoid PAO1) substantially decreased around 12-14, hours while the other *S. aureus* populations survived, suggesting that changes seen in *S. aureus* during this coculture assay are due to killing not growth inhibition.

**S3 Fig. All *S. aureus* isolates grow well when alone.** Average cfu/ml of each *S. aureus* isolate “alone” controls during coculture assay (these exact numbers were used to calculate fold change in Fig 2). You can see little variation among the cfu/ml of the isolates after 24 hours in coculture assay conditions, suggesting that the fold change seen in Fig 2 is not an artefact of differences in *S. aureus* growth.

