## Supplementary figures and images for "Genotypic and phenotypic diversity of *Staphylococcus aureus* from cystic fibrosis lung infections and their interactions with *Pseudomonas aeruginosa*"

### Supplemental Fig 1

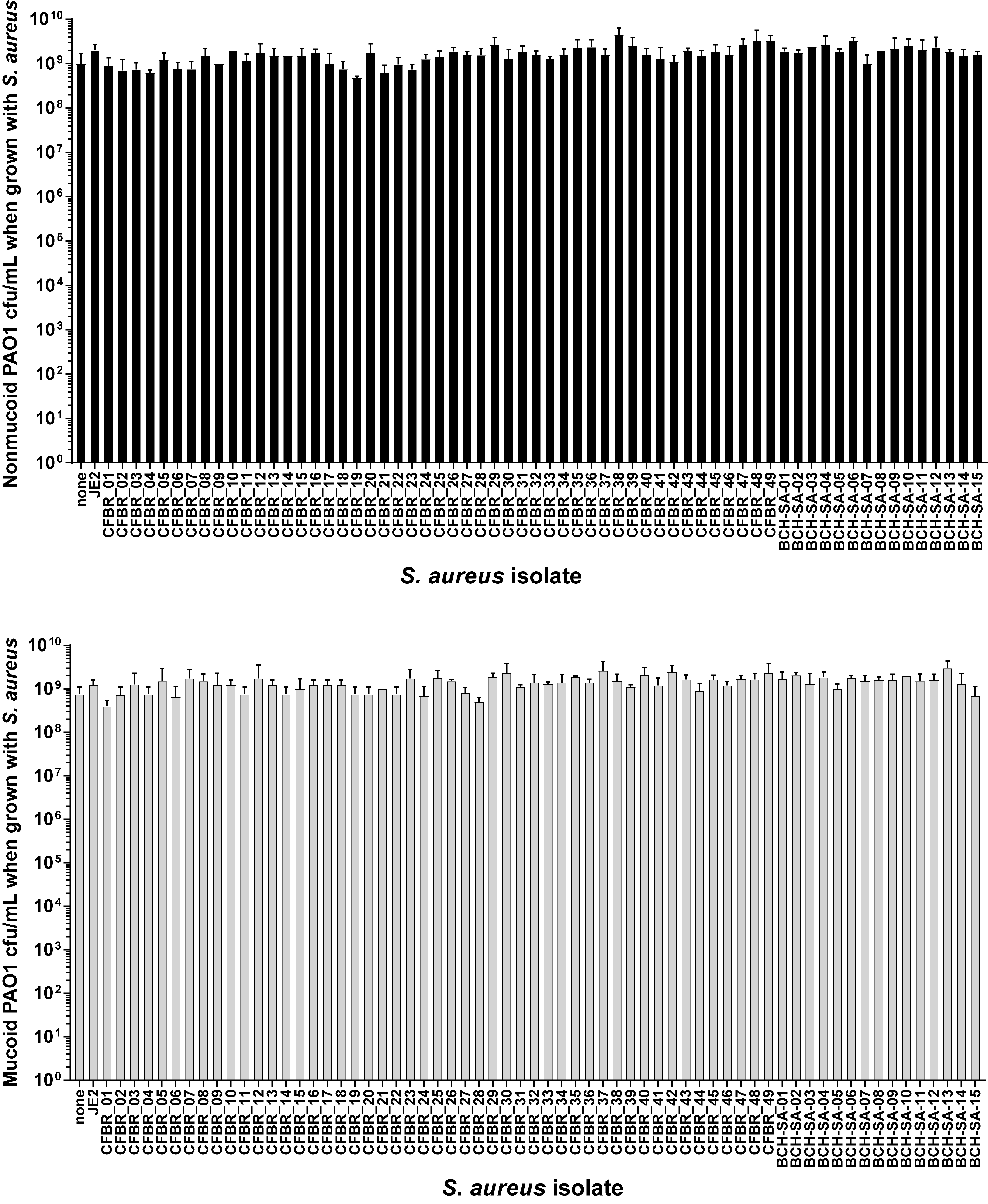

### Supplemental Fig 2

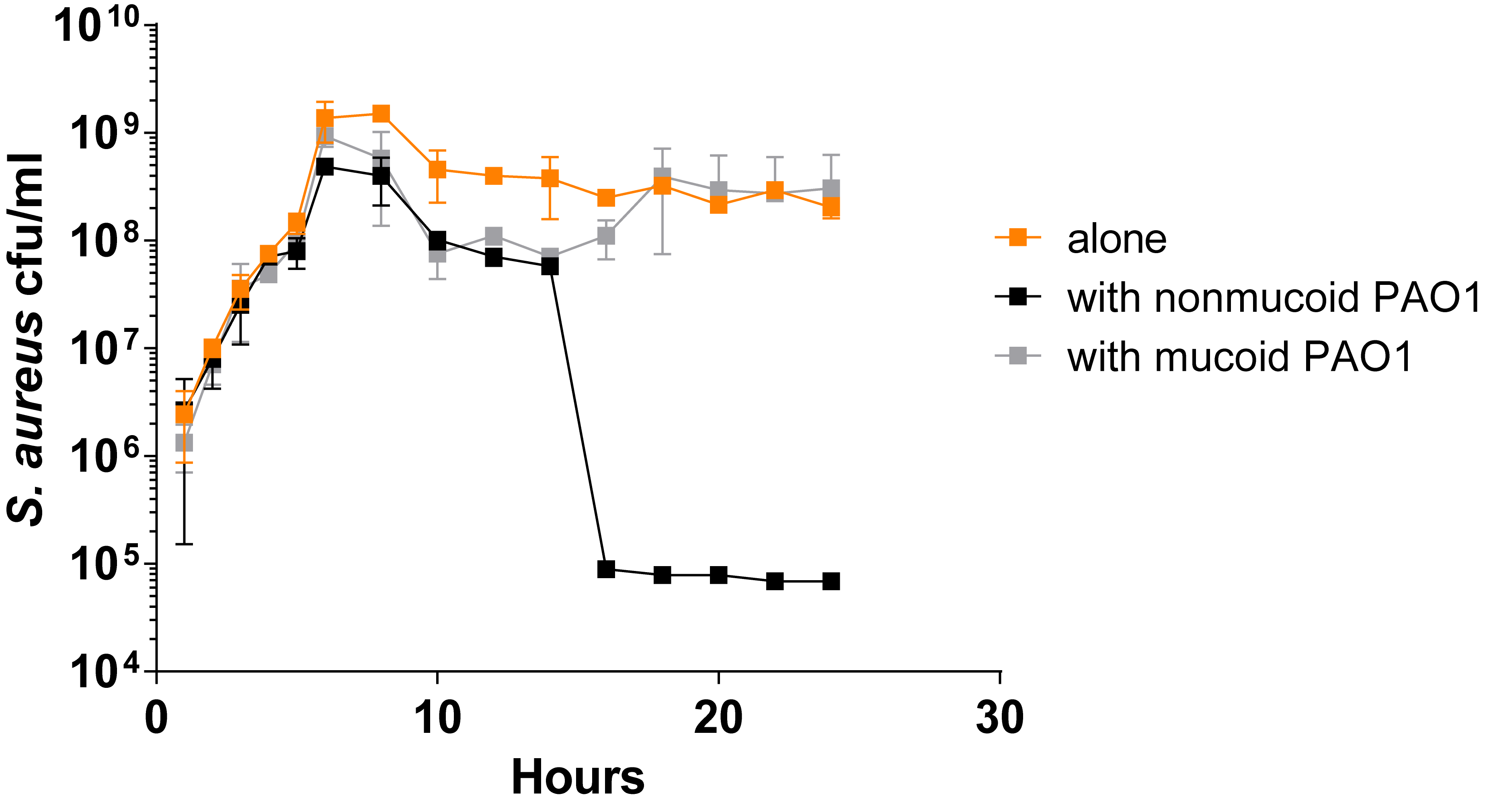

### Supplemental Fig 3

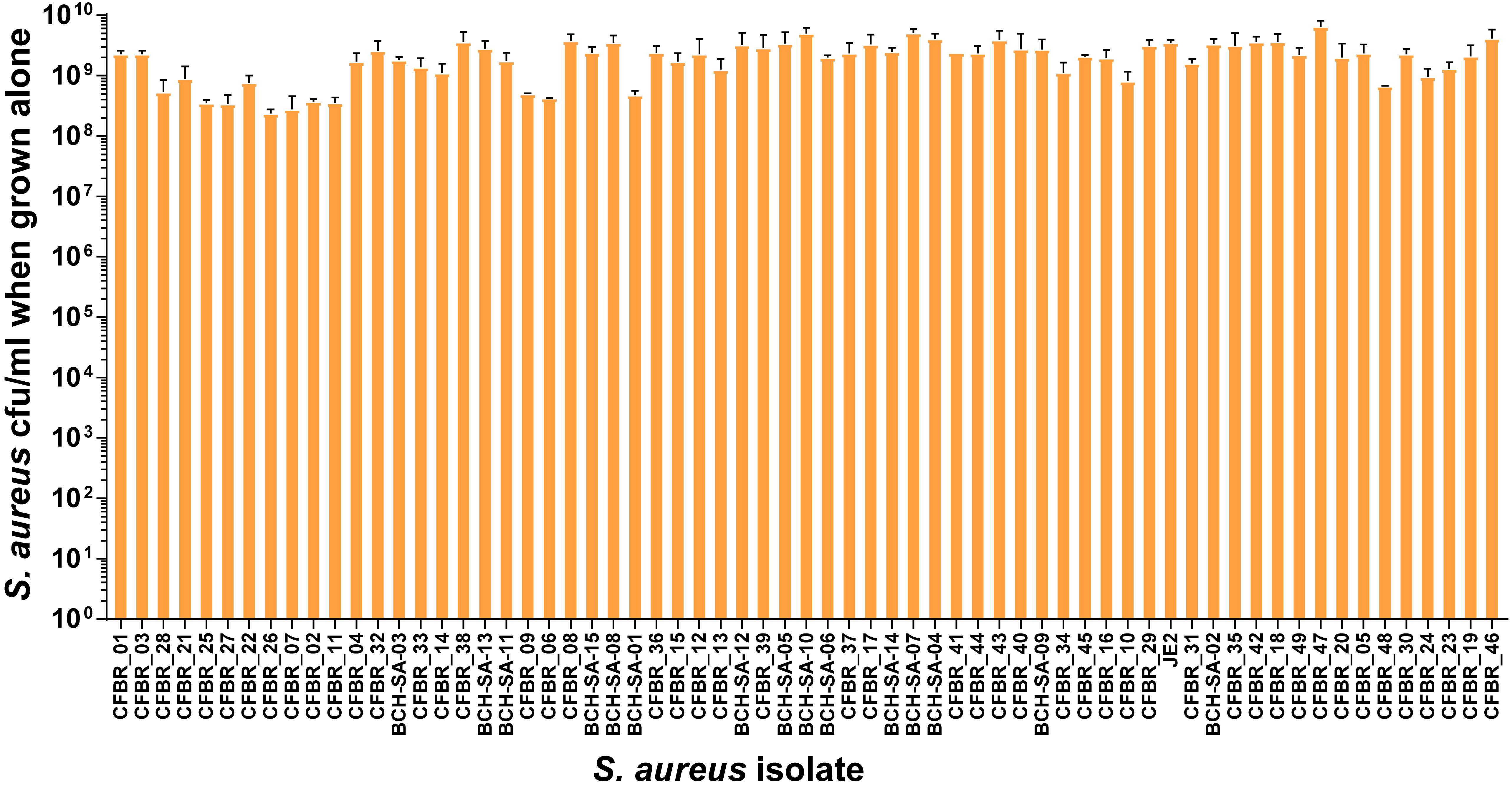
